# *Neuromorpha vorax*: a previously unculturable cosmopolitan protist with an unexpectedly complex life-cycle belonging to Glissomonadida Clade-U/Group-TE

**DOI:** 10.1101/2025.03.13.642991

**Authors:** Gabrielle Corso, Lindsay R. Triplett, Daniel J. Gage

## Abstract

Soil protists are increasingly recognized as common members of complex communities that associate with plant root systems, though their contributions to these communities and to the plant host remain obscure. Members Clade-U/Group-TE, are cosmopolitan soil protists and are often among the most abundant protists associated with plant roots. Here we describe the isolation, culturing and characterization of *Neuromorpha vorax*, a member of the previously uncultured Clade-U/Group-TE branch of the order Glissomonadida. *N. vorax* grew readily when provided bacteriophage lysate as a food source. This allowed us to grow large numbers of the organisms from single-cell isolates and provided ideal conditions for following transitions from one morphology to another. *N. vorax*, like most Glissomonads, has a small, flagellated gliding form, but it also displays a wide range of other morphologies including: a crawling form; small and large trophozoites with multiple, long filopodia; small and large resting cysts; clusters of large dividing cells and it displayed cannibalistic feeding behaviors. Given the small size of most Glissomonads, it may be that other members of this important group, known from environmental surveys, but currently uncultured, might also be readily grown on bacteriophage lysates. In addition, given an abundant food source and clear viewing by microscopy, Glissomonads and other small protists may be found to have life cycles and behaviors that are more complex than is currently appreciated.

**Importance:** Protists from the Clade-U/Group-TE cluster of Glissomonads are wide-spread and abundant colonizers plant roots. Despite being known for over 30 years, they have remained uncultured. We show these protists can be easily cultured using an unusual food source, viral lysates of bacteria. This culturing method allows growth of high numbers of these organisms and reveal thatthey have an unexpectedly complex lifecycle that includes community feeding and cannibalism. Some other, currently unculturable, protists can perhaps be grown with these methods and many of these may also show unexpectedly complex lifecycles. The growth of eukaryotes on virus lysates raises the possibility that viruses in soil may directly contribute to the growth (and not just death) of eukaryotes in soil and root-associated communities.

## Introduction

The Glissomonadida, an order of protists in the phylum Cercozoa, are generally considered bacterivorous and are often among the most abundant protists in terrestrial ecosystems (1–7). They are genetically diverse, but morphologically rather uniform. The Glissomonadida consists of roughly a dozen distinct clades. Of these, no member of clades U, Y, Z and Group-TE have yet to be cultured or characterized beyond sequencing of 18S rRNA genes from environmental gene surveys. Glissomonads are usually small (< 10 µm), bi-flagellate, glide along surfaces using a posterior flagellum, and are considered to have only weak amoeboid characteristics (8–10). There are exceptions to this rule of uniformity. *Proleptomonas faecicola* grows as clusters of dividing cells (11); *Saccharomycomorpha* grows osmotrophically as clusters of dividing cells with no gliding form (12); *Orciraptor* and *Viridiraptor* display a wide range of morphologies including gliding and crawling forms, motile amoeboid forms, several forms capable of cell division, amoeboid forms can dissolve algal cell walls and extract the cytoplasm for food (13). These exceptions suggest that Glissomonads are more diverse in morphology and behavior than currently thought.

Glissomonads designated as belonging to Group-TE have been repeatedly detected in in protist communities from around the world but have resisted efforts to be isolated and cultured. These protists are enriched in rhizosphere samples relative to bulk soil and are often among the most abundant protists in these studies (2, 3, 5–7, 14, 15). As described below, we found that protists designated as members of Group-TE, in databases and in the literature, fall into two well-separated clades: a minority belong to Group-TE as designated by Howe et al. (9, 10), but the majority actually belong to Clade-U as designated by Howe et al.

In this paper we describe the isolation, growth and characterization of *Neuromorpha vorax* a member of the previously uncultured Clade-U (a.k.a. Group-TE) protists. *N. vorax* displays a complex life cycle with gliding and crawling forms, several amoeboid feeding forms, several routes of reproduction, and resting cysts. It also displays communal, cannibalistic, behavior during feeding.

## Results

### Identification, purification and culturing of *N. vorax*

A nested PCR approach was used to identify and track Clade-U organisms as complex protists communities from *Zea mays* rhizosphere samples went through repeated cycles of serial dilution and growth. This eventually resulted in an enrichment culture that contained only a Clade-U Glissomonad and a larger protists belonging to *Paratetramitus* were in the culture, as determined by community sequencing of 18S rRNA gene sequences. Repeated attempts to separate Clade-U protists from *Paratetramitus,* by dilution and single-cell isolations, were unsuccessful, probably because Clade-U protists were separated from their food source. Bacteria and bacterial spores purified from the sample did not allow growth of isolated Clade-U protists when added to their culture medium. Given the small size of the Clade-U cells in the enrichment culture (< 4 µm body length) we attempted to grow them using bacteriophage as a food source. Bacteriophage lysates, made by infecting *Sinorhizobium meliloti* with phage N3, allowed robust growth of the Clade-U protists while, unexpectedly, eliminating *Paratetramitus* and suppressing visible growth of bacteria.

Clade-U protists grew on bacteriophage lysate retained by a 100kD cutoff filter but not on the flow-through. Thus, low molecular weight compounds in the lysate were not sufficient for growth, but proteins, viral particles or aggregates from lysed bacteria larger than 100 kD and smaller than 0.45 μm were sufficient. *N. vorax* also grew on lysates generated by infecting *Mycobacterium smegmatis* with Actinophage “Dulcie” (16, 17). Light and SEM micrographs confirmed that little to no bacteria were present when these organisms were grown in phage lysates. Recent papers document that some protists can consume viruses under laboratory conditions and in natural environments (18–20). However viral lysates have rarely, if ever, been used to isolate and propagate protists species and this technique may allow for the culturing of some recalcitrant protists.

Four pure cultures, identical in morphology and 18S rRNA gene sequence, were independently propagated from single Clade-U cells. One of these subcultures was used in the studies described here and the organism was named *Neuromorpha vorax* for its unusually voracious feeding behavior and the neuron-like morphological appearance of some growth stages.

### Molecular Phylogeny

The 18S rRNA gene sequence of *N. vorax* and a group of 100 18S rRNA genes from Glissomonadida and closely related groups were aligned using 18S rRNA structural information and this was used to generate a maximum-likelihood tree (Fig. 1, Dataset S1). 18S sequences denoted “Group-TE” in Genbank and in the curated PR2 database fell into two, well separated, clades. One contained four sequences belong to the Group-TE clade initially recognized by Howe et al. in papers describing the order Glissomonadida (9, 10). In Fig. 1 this clade clusters, with fairly high confidence (87% bootstrap support), with Allapsidae and *Teretomonas,* as was shown previously (9, 10, 12, 13). *N. vorax* and the other nine “Group-TE” sequences cluster with sequences in Clade-U of the Glissomonadida with high confidence (100% bootstrap support). This cluster groups with a clade containing *Agitata* sp. and related organisms (97% bootstrap support). These are part of a larger clade that includes Clade-Z, Proleptomonadidae and Clade-T/*Saccharomycomorpha* (97% bootstrap support).

**Fig. 1.**
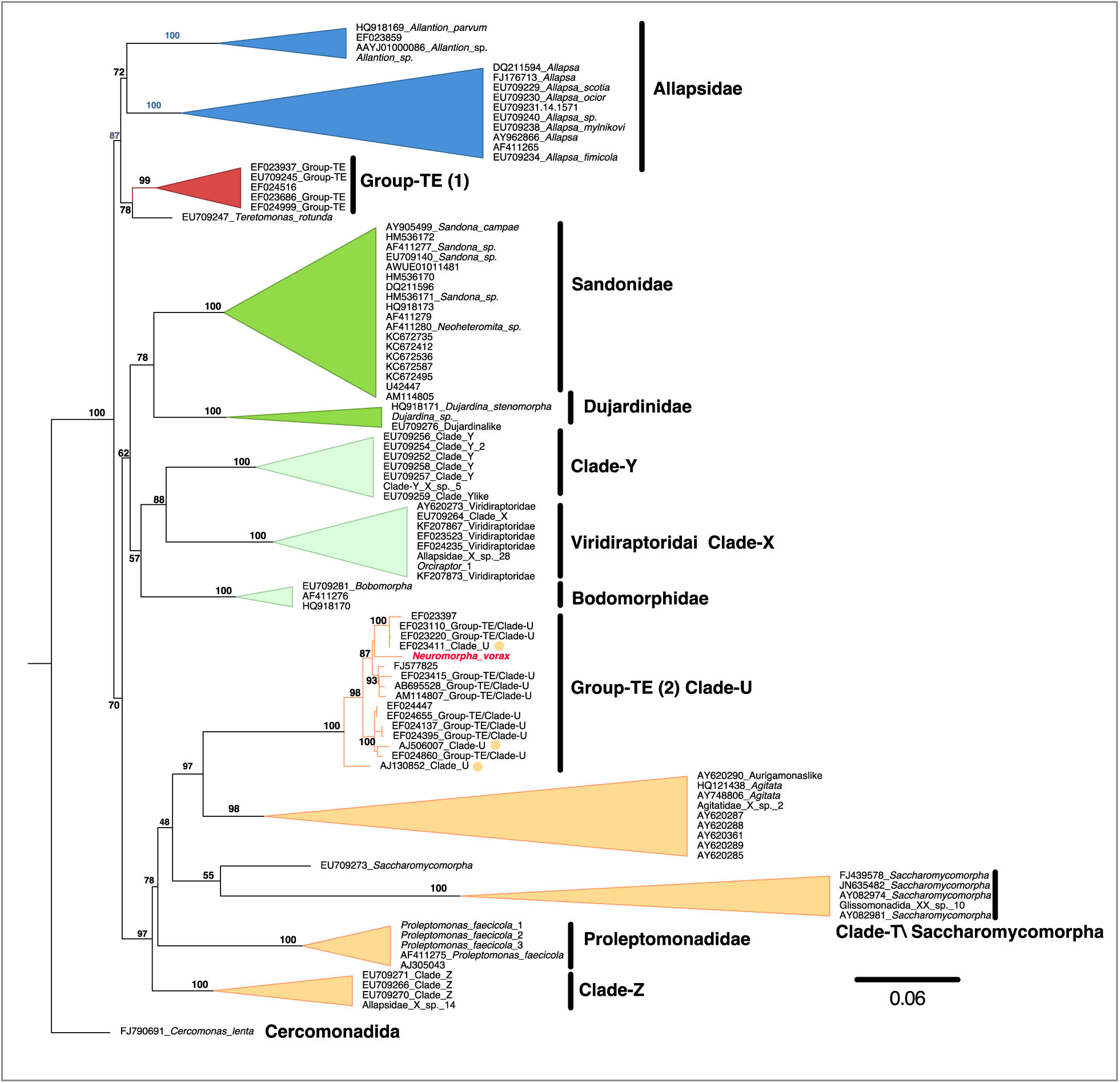
Phylogeny of Glissomonadida and related organisms. Pictured is the best maximum likelihood tree generated by IQ-Tree2, based on a structural alignment of 18S gene sequences, as detailed in Materials and Methods. Sequences identified as Group-TE in the literature but that cluster Clade U sequences are denoted “Group-TE/Clade-U” in the diagram. Support values are real bootstrap values based 100 replicates. A detailed version of this tree is in Supplemental Data Fig 3.

### Clade-U\TE organisms in the rhizosphere

Clade-U ASVs, almost always identified as belonging to “Group-TE” are often some of the most abundant protist ASVs in environmental surveys of soil and they are often enriched in rhizosphere samples relative to nearby bulk soil. Taerum et al. identified two Clade-U (“TE”) ASVs that were enriched in the core protist microbiome of field-grown maize (7). These (P07 and P08) were enriched in the rhizosphere 10- and 30-fold respectively, compared to nearby bulk soil. Clade-U (“TE”) ASVs are also abundant core members of the wheat rhizosphere microbiome samples collected in Europe and Africa, as well as soil-grown lettuce, potato, tomato, and *Arabidopsis* from Germany and the US (5, 6, 15). Simonin et al. identified two Clade-U (“TE”) ASVs in the core rhizosphere microbiome of wheat grown at two sites in Europe and two sites in Africa. One of these was the most abundant and prevalent protist in the core rhizosphere microbiome associated with wheat (6). Sapp et al. (5) identified 5 OTUs (numbers 12,14,106, 291 and 1427) that were rhizosphere enriched in soil-grown *Arabidopsis thaliana* these clustered with Clade-U sequences (Fig. 1 and Dataset S2).

The PR2 v5.0.1 database, which is a curated collection of 18S gene sequences focused on protists, contains 28 sequences identified as belonging to Group-TE (21). 24 of these sequences group with Clade-U and four cluster with Group-TE as defined by Howe et al. (9, 10). Most of the 24 Clade-U sequences derived from soil or were plant-associated (22–25) However, some were isolated from fresh-water aquatic environments such as microbial mats, lake water or drinking water (26–29), indicating that Clade-U may be common in aquatic systems too. These sequences, their associated data and alignments are in Dataset S2 and Fig. S3. Thus, many papers referencing Group-TE as an important member of soil or rhizosphere communities are often referring to members of the Clade-U/*Neuromorpha* cluster.

### Life cycle of *N. vorax*

Culturing *N. vorax* with bacteriophage lysates eliminated visual interference by bacteria or other organisms and revealed a complex life cycle. This included a typical Glissomonad-like gliding form as well as a crawling form, several trophic forms, distinct types of cysts, clusters of dividing cells and communal and cannibalistic behavior. (Fig. 2).

**Fig. 2.**
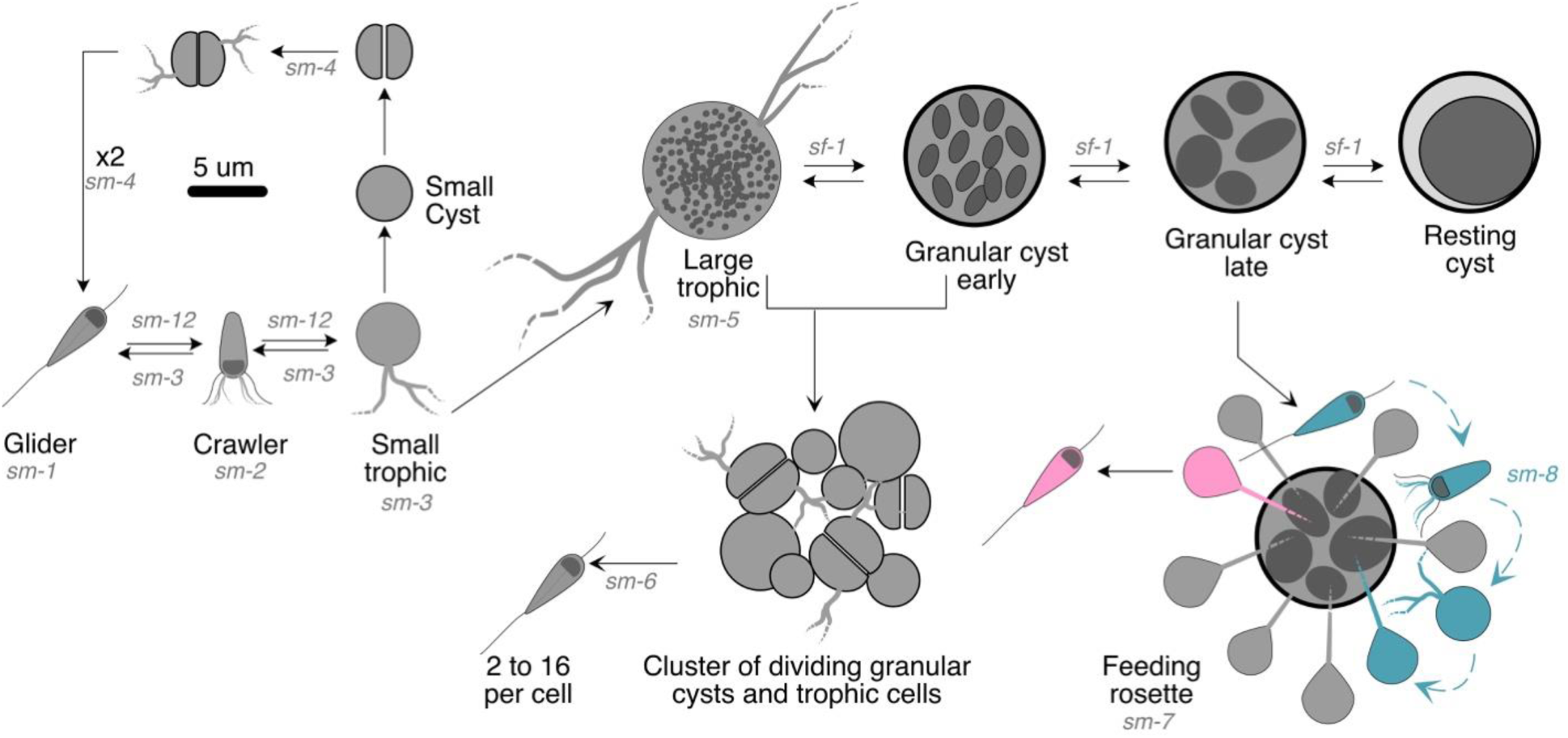
Life cycle of *N. vorax*. Various morphologies and transitions between those morphologies are outlined. Arrows indicate transitions that have been documented by timelapse imaging or still images. Letters on the arrows indicate the supplemental movies (*sm-x*) and supplemental figures (*sf-1*) that document the transitions. The 5μm scale bar shows scale for all the morphotypes in the figure.

### Taxonomy (Neuromorpha vorax)

Phylum Cercozoa Cavalier-Smith 1998

Class Sarcomonadea Cavalier-Smith1993

Order Glissomonadida Howe, Bass, Vickerman Chao and Cavalier-Smith, 2009

### Genus

*Neuromorpha* nov. gen. (urn:lsid:zoobank.org:act:5C12DABA-051D-46C7-95F7-073082FBEFC6).

**Description.** Biflagellate gliders 3-8 µm long. Feeding forms can vary from 5 um to 15 µm and have thin non-anastomosing pseudopods. Can form large (10 µm) resting cysts that adhere to the substrate and are surrounded by a fibrous matrix.

**Etymology:** Genus name refers to the neuron-like appearance of large trophic form.

**Type species:** *Neuromopha vorax*.

**Species:** *Neuromorpha vorax* nov. sp. (urn:lsid:zoobank.org:act:8DCC5AF7-00FA-4BC4-B040-82668A32202E).

**Etymology:** *vorax* because of its voracious feeding habits.

**Collection site:** *Zea mays* root rhizosphere from plants grown at the Connecticut Agricultural Experimental Station Lockwood Farm in July 2021.

**Type material:** Fixed, dehydrated and metal sputtered SEM sample. The hapantotype is the collection of clonal cells grown on a coverslip and prepared as SEM stub3 (Fig. 2H). The hapantotype specimen is deposited in the University of Connecticut Natural History Collections: SEM hapanotype accession number: INV-49193.

**Type sequence:** Genbank accession number: 7661

**Description:** Assumes a variety of forms as described below (Fig. 2, Fig. 3 and Fig. 4).

**Fig. 3.**
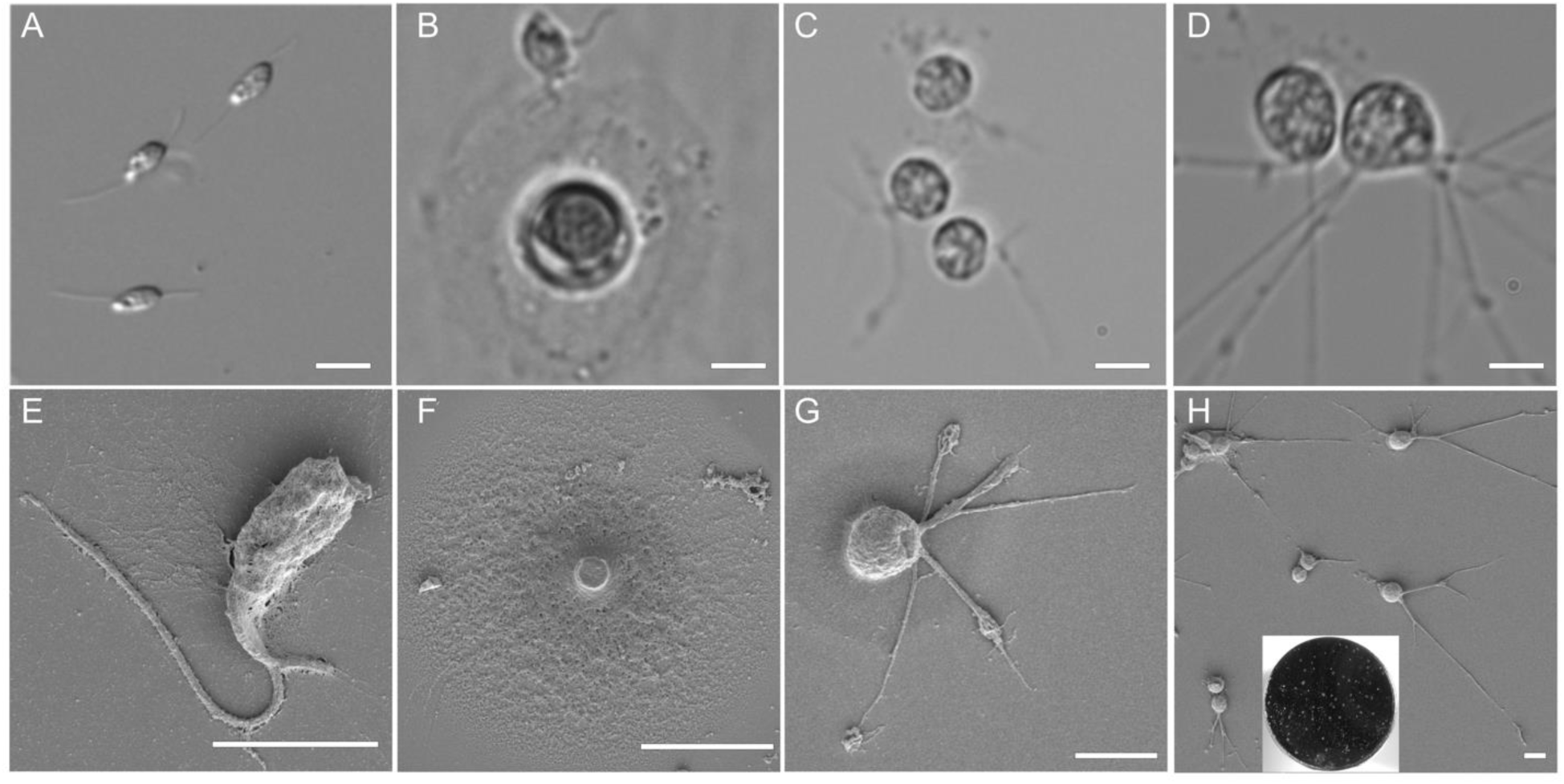
Basic forms of *N. vorax*. A) The biflagellate, gliding, form. B) The crawling (top) form interacting with extracellular material excreted during the formation of a resting cyst (middle). C) Small trophic cells on the bottom of the culture dish. D) a larger trophic cell. D) Large trophic cells on bottom of a culture dish. E) Gliding form transitioning to crawler. F) Large resting cyst. G) Small trophic cell. H) Large, neuron-like, trophic cells. Inset shows the designated hapantotype SEM stub3; white spots are clusters of *N.vorax* of various morphotypes. Panel A is a DIC image, panels B-D are bright-field images, panels E-H are Scanning electron micrographs. Scale bars indicate 5μm. Additional SEM micrographs are in Fig. S2

**Fig. 4.**
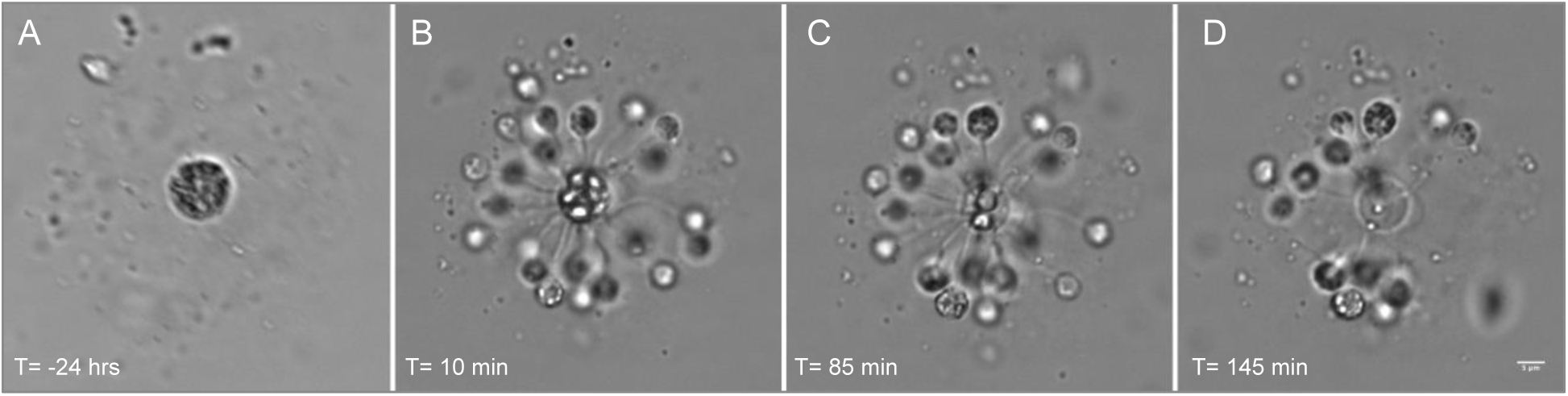
Feeding rosette formed by *N. vorax*. A) The late stage granulated cyst that was fed upon in the feeding rosette. This image was taken ∼24 hours before the rosette itself was imaged. B-D) Feeding rosette consisting of the central cyst and about 25 small trophozoites. Note that the large refractile granules in the central cyst get smaller over time due to feeding and eventually disappear. Time lapse movie of this rosette is Movie S7. Scale of all panels in the same and the scale bar indicates 5μm.

### Gliding form

Gliding forms are biflagellate, cell body is teardrop shaped and 3-9 µm in length, depending on the availability of food. Gliding is smooth with no noted jerking, vibration or other abrupt movements of the cell body. The anterior flagellum is approximately 1.5 times the length of the cell body and pointed forward; the posterior flagellum is approximately 2 times the length of the cell body and trailed behind and is in contact with the substratum during gliding. (Fig. 3A, E, Movie S1).

### Crawling form

Gliding *N. vorax* can slow down, grow pseudopodia and become loosely affixed to the substratum or to films, cysts or aggregates in the growth medium (Fig. 3B). These then use flagella and pseudopods to crawl on the object to which they are attached. Crawlers could then either retract the pseudopods and transition back to gliding, or they would retract their flagella and remain attached to the substrate as small amoeboid trophozoites. (Movie S2).

### Small trophozoites

Small trophozoites have round cell bodies that were 4.6 +/- 0.8 μm in diameter. These have thin, branched filopodia that most often emanate from one spot of the cell body. Filopodia contain enlarged regions that move in both directions along the length of the filament (Fig. 3C, G). These small trophic forms can transition back to gliders (Movie S3), or they can grow and form large, granulated, trophozoites. The small trophozoite can also form a small division cyst that undergoes binary division resulting in two smaller trophozoites that become gliders (Movie S4).

### Large granular trophozoites

Small trophozoites can enlarge in size and become granulated. This occurs most frequently when grown in high lysate concentrations. Large trophozoites often have very long filopodia and resembled neurons (Fig. 3D,H). Granules can be seen moving along the filopodia in both directions (Movie S5). When grown in SES with phage lysate no obvious particles were taken up, or moved, by the filopodia, suggesting that submicroscopic particles are being used as the food source.

### Granular cysts

Granular cysts form when granulated trophozoites retract their filopodia. Internal granules in large trophozoites merge into larger refractile granules. These cyst-like cells typically had four fates: A) they remained unchanged; B) they divided and gave rise 4 or more gliders; C) they grew and divided giving clusters of cells (Movie S6); or D) the granules consolidated into one large granule and the cell become a large resting cyst, with a fried-egg appearance (Fig. S1).

### Large resting cysts

These are 10.0 +/- 2.0 μm in diameter and contain a single large, refractive granule surrounded by a circular matrix of extracellular material (Fig. 3B, F). This form is resistant to desiccation and can become active after rehydration whereupon the large refractile granule separates into smaller granules. At this stage they and appear and behave like granular cysts.

### Small cysts

Small spherical cyst-like cells, which are 4.0 +/- 0.4 μm in diameter, appear in late stage cultures when *N. vorax* is grown with low levels of phage lysate (10^7^ pfu/ml). Upon reactivation, small cysts divide and then develop in gliding forms (Movie S4).

### Feeding rosettes

Feeding rosettes form when gliding cells detect a late stage granular cyst and attach as small trophozoites following a short intermediate stage as crawlers (Fig. 4, Movies S7 and S8). These small trophozoites are attached with a filament 10 to 15 μm long that is likely to be bundled filipodia. Filaments penetrated the outer layer of the cysts and embed in the large granules contained in the cysts. These granules slowly disappear, presumably they are consumed by the attached trophozoites. When the granules in the cyst are gone, the trophozoites convert back to gliders and swim away. The formation of these rosettes is relatively rare, but they were observed in different cultures at different times. Movie S9 shows a related example in which a single small trophozoite with a long filament consumes an aggregate or cyst at the edge of a submerged coverslip. Other cannibalistic behavior is shown in Movie S10 where gliders recognize a broken filopodium from a large trophic cell, then convert to small trophic cells and consume it.

## Discussion

Sequences designated as belonging to the clade “Group-TE” of the Glissomonadida were first describe by Howe et al. (9, 10). Since then, the Group-TE designation has been given to many sequences from environmental 18S surveys. The organisms from which these sequences derived have been noted as being widespread in terrestrial and freshwater ecosystems. They were often enriched, and abundant, in rhizosphere samples from around the world, indicating that they are well adapted to this niche and thus may have impacts, either beneficial or detrimental, to their host plants. Alignment of the 18S rRNA gene of *N. vorax* with those other Glissomonads and related organisms showed that *N. vorax* and most sequences designated as Group-TE, including most of those known to be rhizosphere-enriched, clustered with sequences designated as Clade-U by Howe et al. This clade, and the clade designated “Group-TE” by Howe et al., are clearly distinct in the analyses presented here (Fig. 1) and a visual inspection of the alignments clearly shows that Group-TE and Clade-U (“TE”) have quite different 18S variable regions (Fig. S3 and Dataset S2). These organisms had not been cultured either through targeted efforts (30), or in research programs designed to isolate and characterize members of the Glissomonadida and their close relatives (9, 10, 31, 32). This led to the idea that growth of these organisms perhaps required special conditions or partner organisms for growth.

We found that N. *vorax* was easy to grow and maintain in a standard soil-extract buffer supplemented with bacteriophage lysates from the soil bacteria *S. meliloti or M. smegmatis*. *N. vorax* did not grow on low molecular weight compounds such as proteins, lipids or other small molecules that passed through a filter with a 100 kD, but did grow on phage particles, and larger bacterial debris retained by the filter.

When grown on media containing bacteriophage lysate, *N. vorax* displayed a complex life cycle with cells developing a variety of morphologies and behaviors not previously associated with members of the Glissomonadida. These included large trophozoites with multiple, long filopodia and large resting cysts that were substrate-attached and resembled fried eggs. *N. vorax* displayed communal and cannibalistic feeding behaviors. It formed feeding rosettes which consisted of a large granulated cyst at the center, ringed by small trophozoites that each consumed the contents of the central cyst through a long filament that was likely bundled filopodia. After the contents of the central cyst were drained by the trophozoites they transformed back to gliders and swam away. This type of communal behavior may require signaling between the central cyst and the trophozoites that attached to it. Alternatively, the feeding trophozoites may signal to motile cells which then join the rosette. Related behaviors also suggested signaling between organisms. For example, gliding forms transitioned to crawlers which interacted with the surface of resting cysts, with other crawlers or with pseudopods that broke from, or were ejected by, large trophozoites (Movie S10).

The work outlined here has implications for culturing and charactering other small, hard to grow, protists. Given the small size of gliding forms of many Glissomonads, it is reasonable to suggest that these too may be culturable on phage lysates. This may allow the isolation and culturing of members from groups that have yet to be cultured (Group-TE, Clade-Y and Clade-Z). In addition, given an abundant and readily consumed food source such as bacteriophage lysate, other species of Glissomonads may be found to have life cycles and behaviors that are more complex than is currently appreciated. Hard to culture protists outside the Glissomonadida be easier to culture and characterize using similar methods. Lastly, the work described here shows that the products of viral activity, viral particles and cellular debris, can be used for the growth of eukaryotes. This suggests that viral activity in complex and crowded environments may directly support the growth (and not just death) of fellow community members.

## Methods

### Enrichment of Group-TE protists and detection by nested PCR

*Z. mays* roots, harvested from Connecticut Agricultural Experimental Station Lockwood Farm, in July 2021, were saturated with sterile water and after seven days the water was collected and 100ml samples put into 24-well microtiter plates containing 1ml of SES buffer (SI appendix) and heat-killed *E. coli* strain DH5a cells were added to give a final OD_595_ of 0.005 (7). Following 4 weeks of incubation these wells were tested for the presence of Clade-U protists using a high-sensitivity nested-PCR approach. The first round of PCR amplified near full-length eukaryotic 18S amplicons. This was done with Promega GoTaq Master Mix (Madison, Wisconsin, USA) using (primers EukBR (5-’TGATCCTTCTGCAGGTTCACCTA-3’ and 18SFU (5’-ATGCTTGTCTCAAAGGRYTAAGCCATG-3’). PCR parameters were: initial denaturation at 95^◦^C for 5min followed by 35 cycles of 95^◦^C for 30’’, 60^◦^C for 60” and 72^◦^C for 90’’. The products were gel purified following electrophoresis on a 1% agarose gel. 1-5 ul of the purified products were used in a second round of amplification with Promega GoTaq Master Mix using primers Clade-U_F (5’-CTGRCGAAACTGCTAGCTG 3’) and Clade-U_R (5’-TTGTGTTGCCACAAGAG GCC-3’). PCR parameters were: initial denaturation at 95^◦^C for 5min followed by 35 cycles of 95^◦^C for 30’’, 64^◦^C for 60” and 72^◦^C for 90’’. The products were gel purified using following electrophoresis on a 1% agarose gel and sent to Azenta (Chelmsford, Massachusetts, USA) for Sanger sequencing. Wells that were positive for Clade-U by this test went through a series of 1:10 to 1:100 dilutions into wells with 1ml of fresh SES and heat-killed *E. coli* DH5a at the concentration listed above. Eventually, wells containing only Clade-U and a *Paratetramitus* sp., as determined by community 18S sequencing, were isolated and propagated.

### Purification, growth and maintenance with phage lysates

Dilution and single-cell isolations to separate Clade-U protists from *Paratetramitus* were unsuccessful, probably because the Clade-U protists were removed from their food source. 100ml of a 10^10^ pfu/ml phage lysate made by growing bacteriophage N3 on *Sinorhizobium meliloti* (33) was added to fresh wells made by diluting a 1 month old mixed culture of the Clade-U protists, *Paratetramitus* sp. and various bacteria. These phage lysate additions promoted the growth of Clade-U protists completely suppressed the growth of *Paratetramitus* and the bacteria in the well. Purified lines of Clade-U, generated by isolating single-cells from an initial lysate grown well, were maintained in either 24-well plates containing 1ml of SES plus 10^7^-10^8^ pfu of N3 lysate, or in 25mm^2^ plastic tissue culture flasks containing 5ml of SES plus 5×10^7^ to 5×10^8^ pfu of N3 lysate. Sequences of the full-length 18S rRNA gene of all four lines were identical.

### Phage lysates

*S. meliloti* strain Rm1021 was grown in TY medium with 250 mg/ml streptomycin for 48 hrs. 200ml of these cells were added to 13 x 100mm glass tubes with 100ml of a N3 bacteriophage stock that had been diluted 10^-2^ to 10^-5^ fold in TY. After 20 minutes, 5ml of TY soft-agar (at ∼ 45^◦^C) was added to each tube and the mixture was poured onto room temperature TY plates. Following overnight incubation at 30^◦^C, plates with “lacy” or near-confluent lysis were overlaid with 5ml of SM buffer (SI Appendix) overnight. The SM overlays were pooled (∼20 - 30ml), filtered sterilized with a 0.45mm syringe filter and dialyzed (12-14 kD cutoff) twice for 12 hours against in 500ml of SM buffer. This was then resterilized by filtration as before and stored at 4^◦^C. Lysates typically contained 10^8^-10^9^ pfu/ml

### Microscopy

Protists were grown in 24-well microtiter dishes and imaging was done through the bottom of the wells on a Nikon TE300 inverted microscope with extra-long working distance objectives. Images and time-lapse videos were captured with a Basler acA2040 monochrome CMOS camera (2056 x 1540 pixels, 3.5µm pixel size). The microscope stage, camera and LED illumination were controlled with Micromanager v. 2.0 gamma (34).

### Sequencing

Genomic DNA was isolated from 0.5ml samples of *N. vorax* grown in 1.2ml of SES + N3 (10^8^ pfu/ml) using a MPbio FastDNA Spin Kit for Soil (San Diego, California, USA). Near full-length 18S rDNA was amplified with primers EukBR and 18SFU using NEB Phusion HF Polymerase Master Mix (Ipswich, MA, USA). PCR parameters were: initial denaturation at 96^◦^C for 60’’ followed by 35 cycles of 94^◦^C for 20’’, 68^◦^C for 30” and 72^◦^C for 90’’. Products were run on a 1% agarose gel, purified from the gel and PCR product sequencing performed by Plasmidsaurus using Oxford Nanopore Technology (Eugene, Oregon, USA).

### Phylogenetic analysis

The 18S gene sequence of *N. vorax* was aligned with 100 other near-full length sequences of Glissomonads from the PR2 database and NCBI RefSseq using the software package ssu-align (35, 36). This package aligns and masks alignments based on small subunit RNA structures from eukaryotes, eubacteria and archaea. The software package IQ-Tree2 (37) was used to generate phylogenetic trees using the GTR+R4+F model for the substitution model, state frequency and rate heterogeneity type. Ultrafast bootstrap, parametric aLRT tests and approximate Bayes test were used as indicators of branch support (38, 39). This initial work showed that masking less conserved regions of the alignment had a detrimental effect on tree reproducibility and branch support. The “rf-only” masking which removed only insert columns in the alignment gave the most reliable trees for the clades of interest (Dataset S3). The “rf-only” masked alignment was then processed by IQ-Tree2 using standard bootstrap values. IQ-tree was allowed to select the best substitution model, state frequency and rate heterogeneity type (TIM+F+I+R4) for the “rf-only” alignment. We found that standard bootstrap was a more conservative and reliable indicator of branch support than were the support values used in the faster, initial, tree building trials. Scripts for ssu-align and IQ-tree2 are provided in SI Appendix. The unmasked alignment in fasta and Stockholm formats and the “rf-only” masked alignment, in fasta format, are provided in Dataset S1.

## Data availability

The 18S rRNA gene sequence is deposited under k Genbank accession number: 766. Descriptions of the genus *Neuromorpha* and the species *Neuromorpha vorax* can be found at Zoobank: *Neuromorpha* nov. gen. (urn:lsid:zoobank.org:act:5C12DABA-051D-46C7-95F7-073082FBEFC6) and *Neuromorpha vorax* nov. sp. (urn:lsid:zoobank.org:act:8DCC5AF7-00FA-4BC4-B040-82668A32202E). 18S rRNA gene sequences, alignments and data on bootstrap values vs alignment masking are included in the supplementary material.

## Code availability

Scripts and sequence data used to make the phylogenetic trees are included in the supplementary materials.

## Biological Materials

Dehydrated cysts of *N. vorax* will be provided to researchers at qualified nonprofit institutions upon request.

## Acknowledgements

The authors would like to thank: Vijayalakshmi Venkatramani and the University of Connecticut Electron Microscopy Facility for help with SEM imaging. Dr. Katrina Menard and Dr. Janine Caira for help in depositing the *N. vorax* hapantotype with the University of Connecticut Natural History Collections. Oindrila Mukhopadhyay for help with maintenance of *N. vorax*. This work was supported by AFRI Foundational Program grant from the United States Department of Agriculture-National Institute of Food and Agriculture (USDA-NIFA) to L.R. Triplett, and D.J (grant 2019-67013-24412).

